# A bone fragment-based protocol for molecular analysis of osteocyte-associated transcripts in human bone specimens

**DOI:** 10.64898/2026.05.20.726438

**Authors:** Chiaki Nishizawa, Soju Seki, Emiko Tanaka Isomura, Mari Namikawa, Kazuma Harada, Yusuke Yokota, Tomonao Aikawa, Susumu Tanaka, Toshimi Michigami, Kazuaki Miyagawa

**Affiliations:** Department of Bone and Mineral Research, Research Institute, Osaka Women’s and Children’s Hospital, Izumi, Japan; Department of Oral and Maxillofacial Surgery, Graduate School of Dentistry, The University of Osaka, Suita, Japan; Department of Oral and Maxillofacial Surgery, Graduate School of Biomedical and Health Sciences, Hiroshima University, Hiroshima, 734-0037, Japan

**Author notes:** **Corresponding author** (K.M.). **Related research article** None.

**Keywords:** Osteocyte-associated transcripts, Residual bone fragments, Clinical bone specimens, Sequential collagenase digestion

## Abstract

Osteocytes play a central role in bone remodeling, mineral metabolism, and skeletal homeostasis, but direct molecular analysis of human osteocytes remains technically challenging because they are embedded within the mineralized bone matrix. Surgically obtained human bone specimens provide valuable material for studying human bone biology; however, surface-associated cells, marrow-derived cells, and adherent soft tissues can confound downstream transcript analysis. Here, we describe a bone fragment-based protocol for preparing surgically obtained human bone specimens for molecular analysis of osteocyte-associated transcripts.

The protocol consists of mechanical trimming, mincing into small bone fragments, repeated washing, and five sequential rounds of collagenase digestion to reduce non-osteocytic cellular components associated with the bone surface and marrow spaces. The remaining mineralized bone fragments are then frozen in liquid nitrogen, cryogenically pulverized, and lysed in TRIzol™ reagent for total RNA extraction. Histological validation using residual maxillary bone specimens showed that sequential collagenase digestion markedly reduced adherent soft tissue and extra-matrix nuclei while preserving osteocyte lacunar occupancy.

This protocol provides a practical workflow for bone fragment-based RNA analysis focused on osteocyte-associated transcripts in human bone specimens.

**Specifications table:** 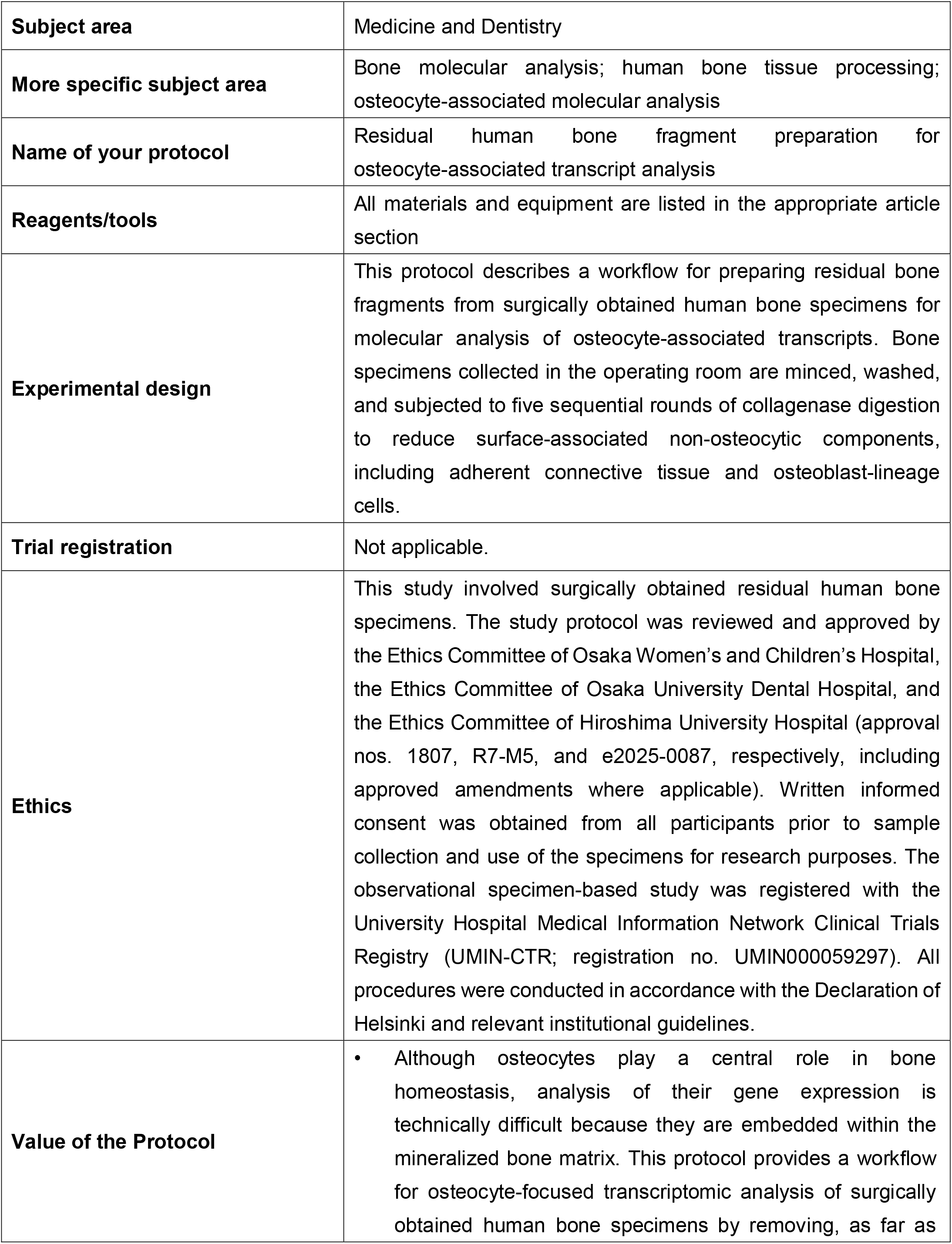

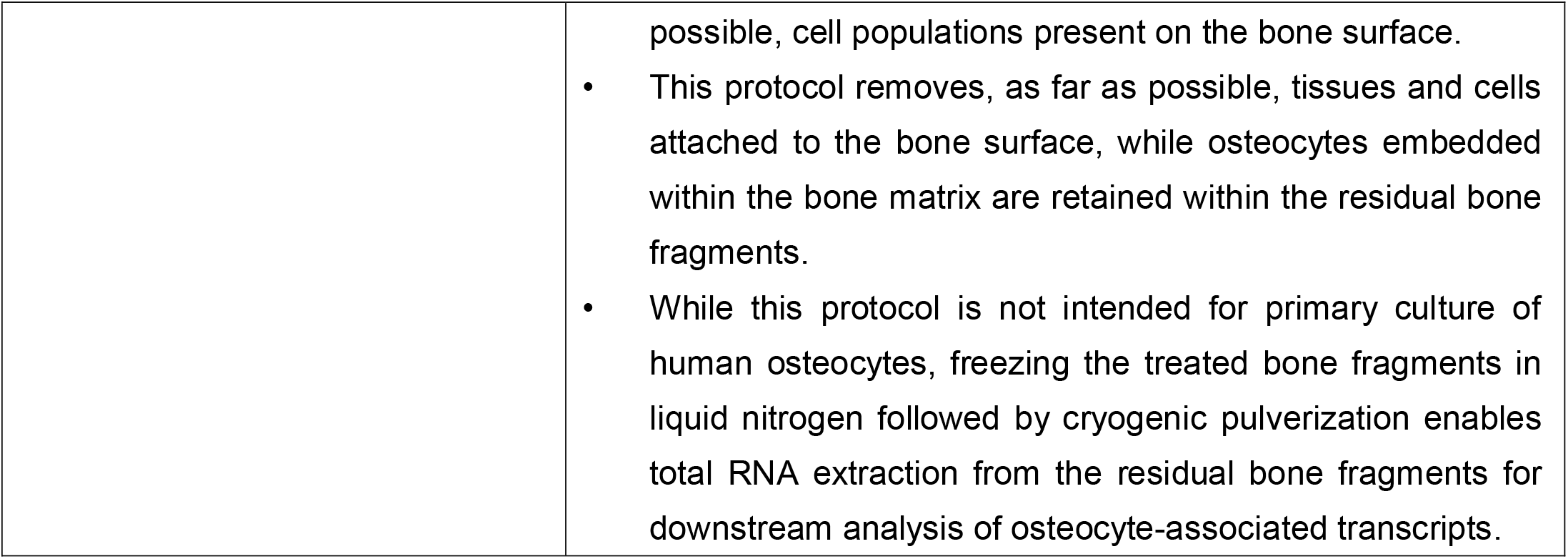

## Background

Osteocytes are terminally differentiated osteoblast-lineage cells embedded within the mineralized bone matrix and represent the most abundant cellular population in bone[1]. Through their extensive cellular network formed by interconnecting dendritic processes, osteocytes act as central regulators of bone remodeling, mineral metabolism, mechanosensation, and skeletal homeostasis[2]. Accumulating evidence also indicates that osteocytes contribute to the pathophysiology of a wide range of skeletal disorders, including metabolic bone diseases[3], rare bone diseases[4,5]. Therefore, molecular analysis of osteocyte-associated signatures in human bone is essential for understanding disease mechanisms that cannot be fully inferred from animal models or *in vitro* osteoblast differentiation systems alone. However, direct molecular analysis of human osteocytes remains technically challenging. Unlike surface-lining osteoblasts, osteoclasts, bone marrow cells, and stromal cells, osteocytes are deeply embedded in a highly mineralized extracellular matrix.

Surgically obtained residual bone specimens provide a valuable opportunity to investigate human bone biology under clinically relevant conditions. Bone fragments collected during oral and maxillofacial or orthopedic procedures may retain information on tissue-specific cellular states, donor-dependent variation, and disease-associated molecular signatures. Nevertheless, a practical and reproducible protocol for preparing small human bone fragments for osteocyte-associated molecular analysis has not been fully standardized.

Here, we describe a bone fragment-based protocol for molecular analysis of osteocyte-associated transcripts in surgically obtained human bone specimens, adapted from murine sequential bone digestion workflows[6,7]. This protocol is designed to reduce cellular contributions from bone surface-associated and marrow-derived populations through mechanical trimming, washing, and sequential enzymatic digestion, while retaining residual mineralized bone fragments for downstream RNA-based analysis. Although this protocol does not isolate human osteocytes as single cells, it provides a practical workflow for preparing bone fragment lysates that can be used to assess osteocyte-associated transcripts in small clinical bone specimens. This workflow may support clinically oriented molecular assessment of residual bone specimens obtained during routine surgical procedures, providing a framework for evaluating osteocyte-associated changes alongside pathological and diagnostic evaluation of bone tissues.

### Description of protocol

#### Materials (Fig. 1)

- Timer
- Tissue forceps
- 50-mL conical tubes
- Petri dishes, 90 mm
- Laboratory sealing film
- 100-mL glass bottle
- 100-mL amber glass bottle
- Mayo scissors, curved blades, ring handle
- Metzenbaum scissors, curved blades, ring handle
- Bone-cutting rongeur
- Stainless steel micro spatula
- Disposable serological pipettes
- Autoclavable silicone pipette bulb
- Sterile disposable plastic transfer pipettes
- Cell strainer, 100-µm mesh
- 1.5-mL microcentrifuge tubes
- Liquid nitrogen Dewar flask

**Fig 1.**
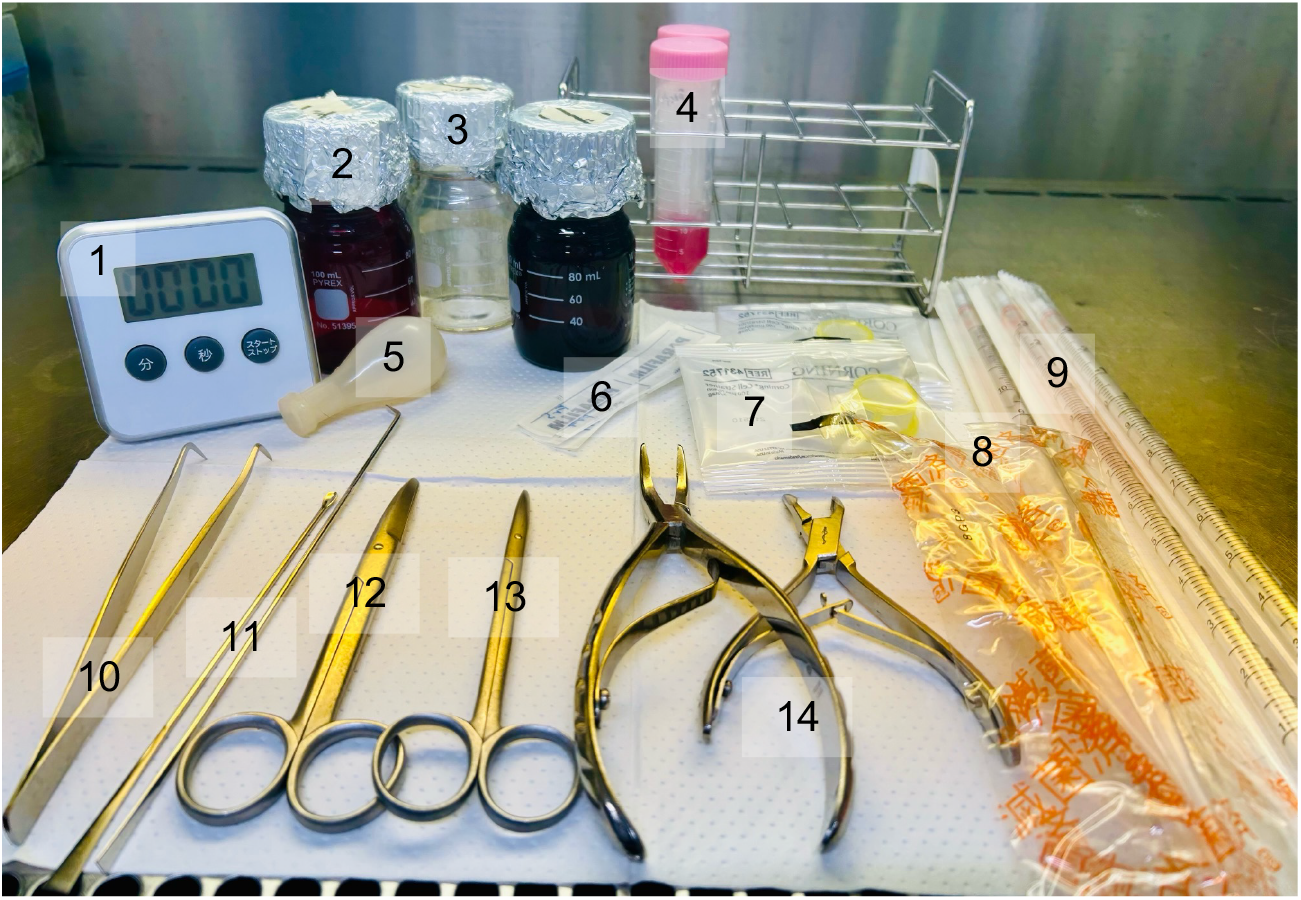
Materials prepared inside the clean bench before starting the experiment. All materials were maintained under sterile conditions. 1, timer; 2, 100-mL amber glass bottle; 3, 100-mL glass bottle; 4, 50-mL conical tube; 5, autoclavable silicone pipette bulb; 6, laboratory sealing film; 7, cell strainer; 8, disposable serological pipettes; 9, sterile disposable plastic transfer pipettes; 10, tissue forceps; 11, stainless steel micro spatula; 12, Mayo scissors; 13, Metzenbaum scissors; 14, bone-cutting rongeur.

### Reagents

- Preparation medium: α-MEM supplemented with penicillin/streptomycin without FBS
- Trimming medium: preparation medium supplemented with 10% FBS; 10 mL per 100-mm dish, 3 dishes, pre-warmed to 37°C
- Trimming PBS(−): PBS(−) supplemented with 10% FBS
- Culture medium: α-MEM supplemented with 10% FBS and 0.25 mM ascorbic acid, stored in a 100-mL amber glass bottle protected from light
- PBS(−): phosphate-buffered saline without Ca^2+^ and Mg^2+^
- Liquid nitrogen
- TRIzol™ reagent (Thermo Fisher Scientific, Waltham, MA, USA; Cat. No. 15596026)
- Collagenase solution: 1.25 mg/mL collagenase in α-MEM supplemented with penicillin/streptomycin, stored in a 100-mL amber glass bottle protected from light Collagenase for cell dispersion, from *Clostridium histolyticum* (FUJIFILM Wako Pure Chemical Corporation, Osaka, Japan; Cat. No. 034-22363) **Note:** Collagenase activity and accompanying protease activities vary among manufacturers, product types, and lots. For substitution, crude collagenase preparations from Clostridium histolyticum with an activity of approximately 100–200 units/mg are recommended as a starting point. Users should perform preliminary optimization of enzyme concentration, digestion time, and fragment size before processing valuable clinical specimens.

### Equipment

- Swing-bucket centrifuge
- CO_2_ incubator
- Temperature-controlled shaking water bath equipped with a stainless-steel spring rack
- Cryogenic tissue pulverizer, e.g., Cryo-Press CP-100W, Sansyo, Japan
- Bone mill, optional, e.g., Bone Mill ex-2, YDM, Japan

### Procedure

#### Step 1 - Acquisition and initial preparation of human bone specimens

1. Collect residual human bone specimens obtained during surgery aseptically and immediately place them into 50-mL conical tubes containing preparation medium pre-cooled to 4°C. **Note:** Do not bring serum-containing culture medium into the operating room. Serum-derived products may pose a theoretical risk of introducing unknown pathogens or biological contaminants into the surgical environment. Therefore, human bone specimens should be collected intraoperatively only into serum-free preparation medium, and any serum-containing medium should be introduced only after transfer to the laboratory.
2. Transport the specimens to the laboratory on ice within 30 min.
3. In a clean bench, open the tube cap and transfer the bone specimen to a 90-mm Petri dish containing trimming PBS(−) (Fig. 2A). **Note:** During initial trimming, the bone specimen should be kept in a serum-containing or blood component-containing solution. Trimming PBS(−), trimming medium, or preparation medium containing blood-derived components from the collected specimen may be used according to the operator’s preference.
4. Gently remove adherent soft tissues from the specimen using two pairs of tissue forceps.
5. Optional: For bone specimens that are too hard to cut with Mayo scissors, process the specimen into small particles using a bone mill before subsequent handling (Fig. 2B). **Note:** Preliminary fragmentation with a bone mill may not be feasible for large or dense bone specimens. When necessary, the specimen should be reduced intraoperatively to a size suitable for milling using an irrigated ultrasonic cutting device or manual bone-cutting instruments, such as bone forceps or rongeurs. Highly invasive rotary cutting instruments should be avoided because they may cause excessive heat generation or mechanical damage, potentially affecting downstream RNA-based analyses.

**Fig 2.**
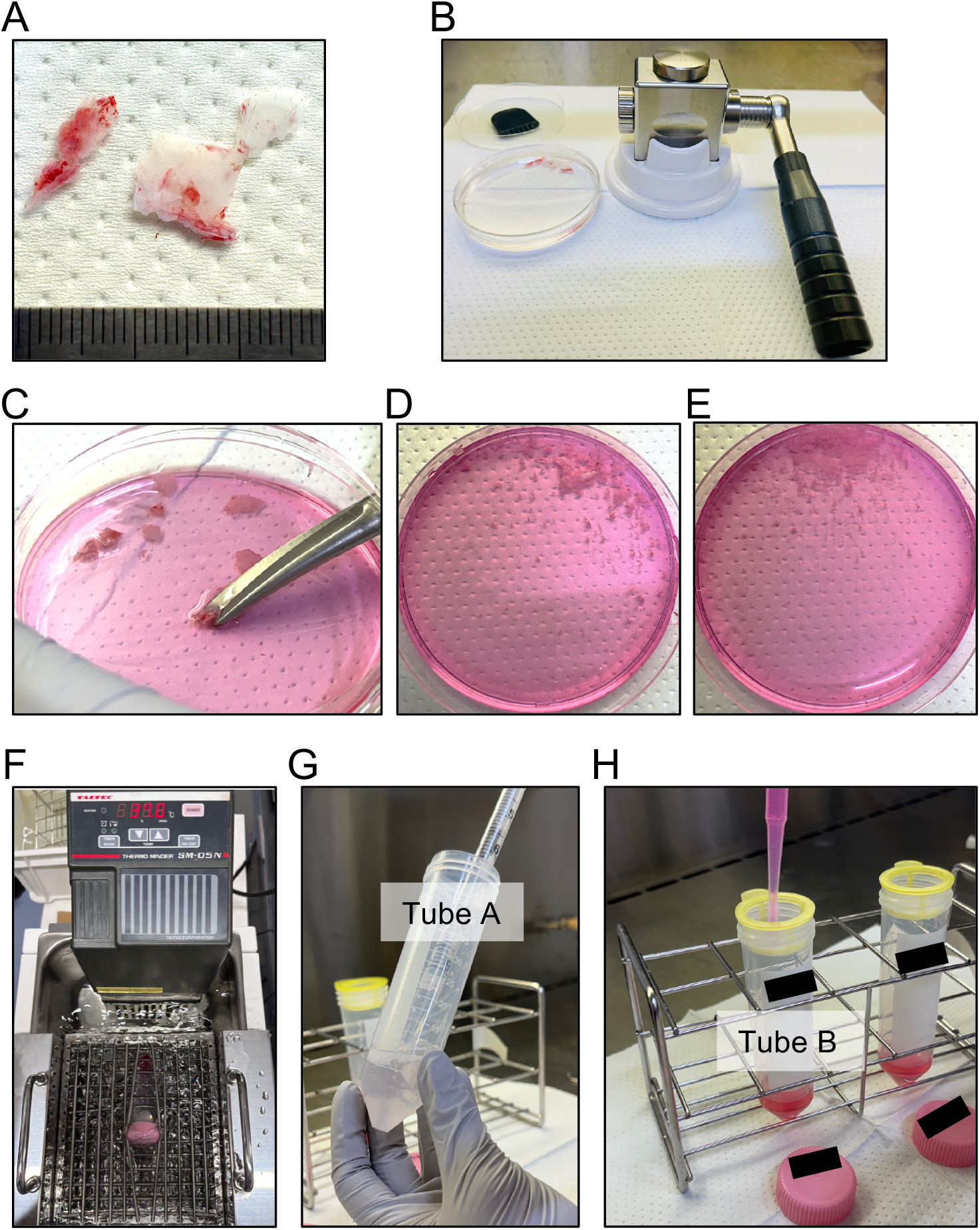
Workflow for sample preparation and enzymatic digestion of bone fragments. A, bone fragments obtained from the maxillary sinus wall before mincing; B, preliminary fragmentation using a bone mill; C, mincing of bone fragments using Mayo scissors; D, bone fragments after mincing with Mayo scissors; E, bone fragments after subsequent mincing with Metzenbaum scissors; F, collagenase digestion in a shaking water bath; G, pipetting of the suspension in tube A after enzymatic digestion; H, filtration of the supernatant through a 100-µm cell strainer into tube B.

#### Step 2 – Sample preparation

1. Mince the bone specimens into small fragments, approximately 0.5–1.0 mm in size, using sterile Mayo scissors for 5–15 min, followed by sterile Metzenbaum scissors for another 5–15 min (Fig. 2C). Care should be taken to obtain fragments of relatively uniform size (Fig. 2D, E). **Note:** Bone specimens should be minced with Mayo scissors and then Metzenbaum scissors until the fragments can be cut with minimal resistance. Uniform fragment size is critical for reproducible collagenase digestion, as larger fragments may be insufficiently digested. Because bone fragments can easily scatter during mincing, the Petri dish should be partially covered with its lid while cutting.
2. Using a sterile spatula, transfer the minced bone fragments into a 50-mL conical tube containing 10 mL of FBS-free preparation medium.
3. Centrifuge the tube at 1,000 rpm for 30 s using a swing-bucket centrifuge, and carefully discard the supernatant.
4. Add 10 mL of FBS-free preparation medium to the tube. Gently invert the tube 5–10 times to wash the bone fragments.
5. Centrifuge the tube again at 1,000 rpm for 30 s, and carefully discard the supernatant.
6. Repeat the washing procedure twice more to further remove residual blood, bone marrow components, and loosely attached cells.

#### Step 3 – Enzymatic digestion of bone fragments

1. Add 10 mL of collagenase solution to the 50-mL conical tube containing the washed bone fragments, hereafter designated tube A. Seal the gap between the tube and cap with laboratory sealing film. Place tube A horizontally in a shaking water bath and incubate with shaking at 37°C for 15 min (Fig. 2F). **Note:** The shaking speed should be adjusted so that the bone fragments can move gently within the collagenase solution. Excessive shaking should be avoided, as it may increase mechanical stress on the released cells.
2. After incubation, pipette the suspension up and down for 2 min using a 10-mL disposable serological pipette and a silicone pipette bulb to detach cells from the bone fragments (Fig. 2G). **Note:** During pipetting, avoid excessive foaming, as foam formation can impose unnecessary mechanical stress on the released cells and may reduce cell quality. Pipette the suspension gently and steadily, keeping the pipette tip immersed in the liquid whenever possible.
3. Allow tube A to stand for 30 s to let the bone fragments settle. Transfer the supernatant through a 100-µm cell strainer into a fresh 50-mL conical tube containing 10 mL of culture medium, hereafter designated tube B (Fig. 2H). **Note:** The supernatant containing released cells is transferred into serum-containing culture medium to terminate or attenuate collagenase activity.
4. Add 10 mL of PBS(−) to the remaining bone fragments in tube A, and pipette the suspension up and down for 2 min (Fig. 2G).
5. Allow tube A to stand for 30 s, then transfer the supernatant through the 100-µm cell strainer into tube B.
6. Centrifuge tube B at 1,000 rpm, approximately 160 × g, for 5 min using a swing-bucket centrifuge.
7. Carefully remove the supernatant and resuspend the cell pellet in 2 mL of fresh culture medium.
8. Keep the cell suspension in tube B at 37°C in a CO_2_ incubator. This cell suspension is designated Fraction 1.
9. Add 10 mL of fresh collagenase solution to the remaining bone fragments in tube A.
10. Repeat the digestion, pipetting, filtration, centrifugation, and resuspension procedures described above four additional times to obtain Fractions 2–5. **Note:** This protocol was designed for residual bone specimens of up to approximately 0.5 g per 50-mL conical tube. Because collagenase digestion efficiency may vary depending on bone type, specimen size, and the amount of bone tissue processed, larger specimens may require division into multiple tubes and additional optimization of collagenase concentration, digestion time, and fragment size.

#### Step 4 – Preparation of TRIzol lysates from fractionated cells and residual bone fragments

1. Discard the cell suspensions obtained from Fractions 1 and 2. **Note:** Fractions 1 and 2 are not used for the osteoblast-enriched RNA fraction in this protocol, because these early fractions are expected to contain a higher proportion of loosely attached cells, blood-derived cells, and bone marrow-associated cells.
2. Combine the cell suspensions obtained from Fractions 3-5 into a pooled fraction, designated the osteoblast-enriched fraction, according to previously described mouse bone osteoblast/osteocyte fractionation methods [6,7]. **Note:** This pooled fraction is expected to be enriched for bone surface-associated osteoblast-lineage cells relative to the early digestion fractions and can be processed as a comparative cellular fraction when needed.
3. Centrifuge the osteoblast-enriched fraction at 1,300 rpm for 5 min using a swing-bucket centrifuge.
4. Carefully discard the supernatant and resuspend the cell pellet in 1 mL of TRIzol™ reagent.
5. Store the TRIzol™ lysate at −80°C until total RNA extraction.
6. Before freezing, remove excess liquid from between the residual bone fragments as much as possible using a twisted Kimwipe or other absorbent laboratory tissue (Fig. 3A).
7. Freeze the residual bone fragments in liquid nitrogen and pulverize them using a cryogenic tissue pulverizer (Fig. 3B, C).
8. Quickly transfer the pulverized bone powder into 1 mL of TRIzol™ reagent (Fig. 3D).
9. Store the bone fragment lysate at −80°C until total RNA extraction. **Note:** This protocol describes the preparation of fractionated cell and bone fragment lysates in TRIzol™ reagent. Subsequent total RNA purification can be performed using standard TRIzol-based extraction procedures or column-based cleanup methods according to the user’s downstream application.

**Fig 3.**
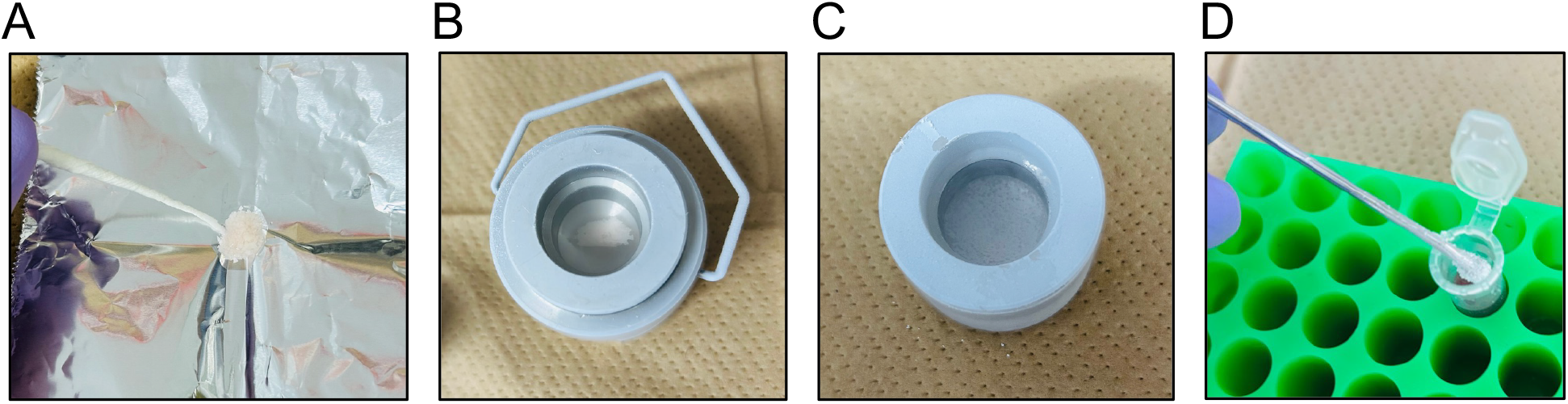
Preparation of residual bone fragments for cryogenic pulverization. A, excess liquid is removed from residual bone fragments using a twisted absorbent laboratory tissue; B, residual bone fragments are frozen in liquid nitrogen before pulverization; C, bone powder after cryogenic pulverization; D, pulverized bone powder is transferred into a 1.5-mL tube containing TRIzol™ reagent.

## Protocol validation

Protocol validation was performed using residual maxillary bone specimens collected from three patients who underwent Le Fort I osteotomy as part of orthognathic surgery. The specimens were obtained from the posterior and lateral wall regions of the maxillary sinus.

### 1. Changes in bone fragments due to sequential collagenase digestion

Representative macroscopic images show that minced bone fragments were initially mixed with red blood cells and marrow tissue (Fig. 4A). Three repeated washes by PBS(−) reduced suspended blood and marrow-derived components (Fig. 4B), and sequential collagenase digestion further reduced visible marrow tissue associated with the residual bone fragments (Fig. 4C).

**Fig 4.**
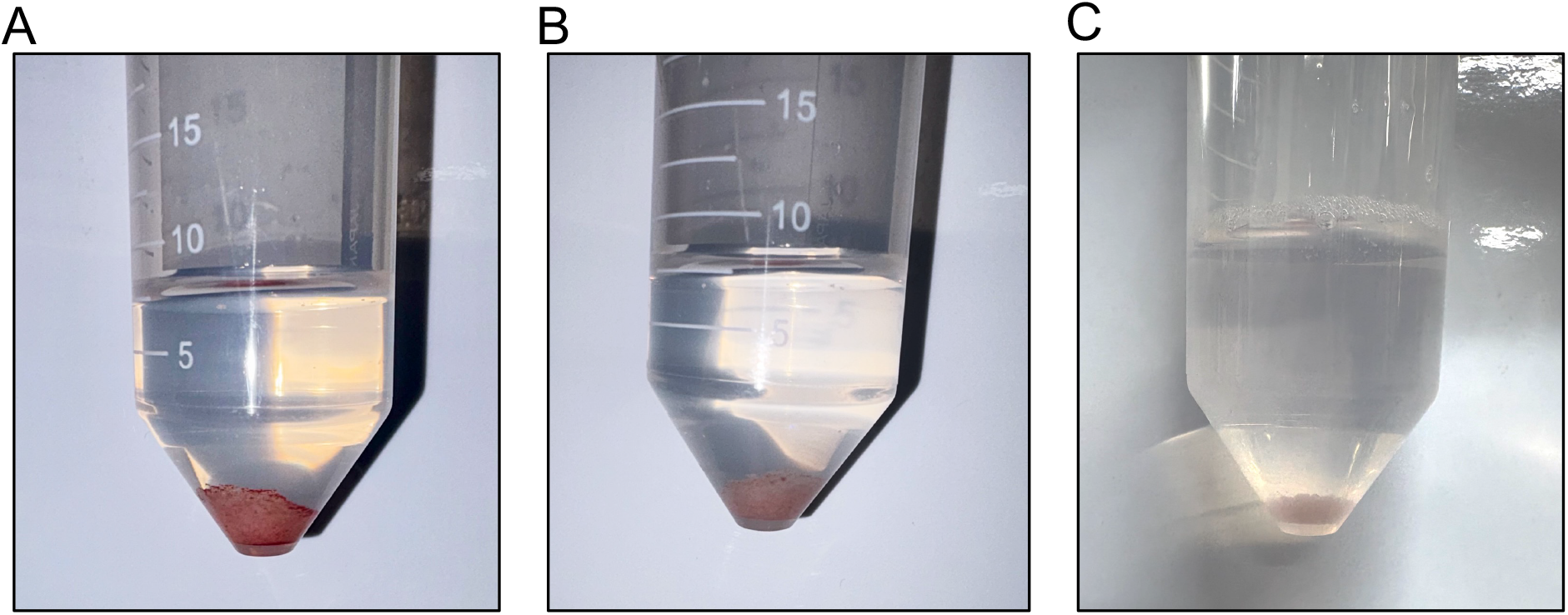
Representative macroscopic changes in bone fragments before and after processing. A, bone fragments immediately after mincing; B, bone fragments after three repeated washes by PBS(−); C, residual bone fragments after five rounds of collagenase digestion.

### 2. Histological validation of surface-associated cell removal by sequential collagenase digestion

H&E-stained sections of bone fragments before and after sequential collagenase digestion were analyzed. Post-digestion fragments showed minimal residual bone marrow tissue, as shown in Fig. 5A. For each sample, five fields were randomly selected from low-magnification images and evaluated at high magnification using a ×20 objective. The mean value of the five fields was used as the representative value for each sample. Soft-tissue coverage of bone fragments (STC) and the extra-matrix nuclei ratio (EMNR) were significantly decreased in post-digestion fragments compared with pre-digestion fragments, reaching nearly negligible levels (*P* = 0.036 for STC; *P* = 0.003 for EMNR; Fig. 5B, C). In contrast, osteocyte lacunar occupancy (OLO) was comparable between pre-digestion and post-digestion fragments (*P* = 0.939; Fig. 5D).

**Fig 5.**
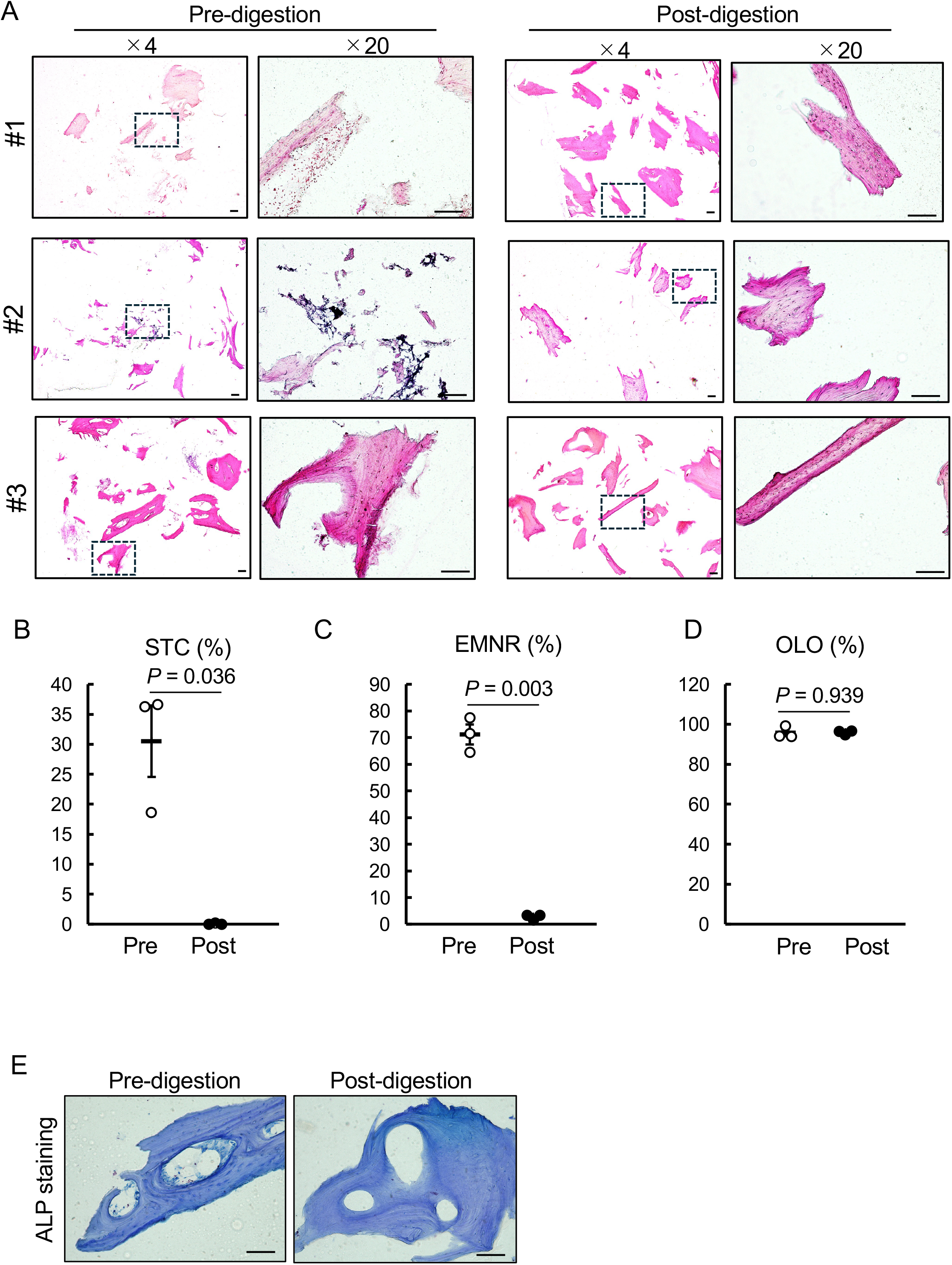
Histological validation of bone fragments before and after sequential collagenase digestion. A and E, representative decalcified frozen sections of bone fragments before and after digestion. Section thickness was 14 μm. A, H&E-stained sections of bone fragments from three independent specimens before and after digestion. Low-magnification images were acquired at ×4, and boxed areas are shown at ×20. B–D, quantitative analysis of soft-tissue coverage of bone fragments (STC; B), extra-matrix nuclei ratio (EMNR; C), and osteocyte lacunar occupancy (OLO; D). For quantification, five ×20 fields containing bone fragments were selected using systematic random sampling for each sample before and after digestion, and the mean value was used as the representative value for each sample. STC was defined as the percentage of bone surface length covered by adherent soft tissue or cellular layers relative to the total bone surface length. EMNR was defined as the percentage of nuclei located outside the mineralized bone matrix among total nuclei. OLO was defined as the percentage of osteocyte lacunae containing nuclei among total osteocyte lacunae. Data are presented as individual data points with mean ± SEM. Statistical comparisons were performed using Welch’s t-test. E, ALP-stained sections of bone fragments before and after digestion using a commercial staining kit (FUJIFILM Wako Pure Chemical Corporation, Osaka, Japan; Cat. No. 294-67001). Scale bars, 100 μm.

To further assess the reduction of surface-associated cellular components, including osteoblast-lineage and stromal cell populations, ALP staining was performed on bone fragment sections before and after sequential collagenase digestion (Fig. 5E). ALP-positive cells on the bone surface were rarely detected in post-digestion fragments. Furthermore, soft tissue components within intracortical cavities, presumed to correspond to Haversian canals, were largely removed after sequential collagenase digestion.

These findings indicate that sequential collagenase digestion efficiently reduces surface-associated cells and adherent soft tissue surrounding the bone fragments while preserving osteocyte-containing lacunae within the mineralized bone matrix.

## Limitations

This protocol reduces surface-associated cellular components from surgically obtained residual bone specimens by sequential collagenase digestion and enables molecular analysis focused on osteocyte-associated transcripts. However, the resulting residual bone fragments do not represent a completely pure osteocyte population or an osteocyte-specific compartment. RNA extracted from these fragments may still contain signals from non-osteocyte components, and the results should therefore be interpreted together with complementary analyses, such as histological assessment, immunostaining, or *in situ* hybridization.

The proportion of residual surface-associated cells may vary depending on specimen size, anatomical site, tissue architecture, and pathological condition. Therefore, a small portion of the processed bone fragments should be examined histologically for each sample whenever possible. This quality-control step may help interpret sample-to-sample variability in downstream gene expression analyses.

Further, the present workflow is intended to prepare residual bone fragment lysates for RNA-based analysis rather than to isolate or culture viable human osteocytes. Therefore, this protocol should not be directly applied to osteocyte culture without substantial modification and validation. Unlike previously reported protocols aimed at viable osteocyte isolation from human bone[8], this approach does not rely on chelation-mediated decalcification for osteocyte release and instead directly processes residual mineralized bone fragments for total RNA extraction.

## CRediT author statement

**Chiaki Nishizawa:** Methodology, Validation, Formal analysis, Investigation, Data curation, Writing – review & editing, Visualization, Funding acquisition. **Soju Seki:** Investigation, Resources, Writing – review & editing. **Emiko Tanaka Isomura:** Investigation, Resources. **Mari Namikawa:** Resources. **Kazuma Harada:** Resources. **Yusuke Yokota:** Resources. **Tomonao Aikawa:** Project administration, Resources. **Susumu Tanaka:** Project administration, Resources. **Toshimi Michigami:** Supervision, Writing – review & editing. **Kazuaki Miyagawa:** Conceptualization, Methodology, Validation, Formal analysis, Data curation, Writing – original draft, Writing – review & editing, Visualization, Supervision, Project administration, Funding acquisition.

## Acknowledgments

This study was supported by Grants-in-Aid for Scientific Research from the Japan Society for the Promotion of Science (JSPS KAKENHI; grant numbers 25K24084 to C.N., and 22K10216 and 25K02829 to K.M.). Additional support was provided by the Osaka Medical Research Foundation for Intractable Diseases to C.N., and by the Open and Transdisciplinary Research Initiatives (OTRI), Osaka University, the Nakatomi Foundation, and the Mother and Child Health Foundation to K.M.

We thank Editage (http://www.editage.jp) for the English language editing.

## Declaration of interests

☒ The authors declare that they have no known competing financial interests or personal relationships that could have appeared to influence the work reported in this paper.

## Notes

### Competing Interest Statement

The authors have declared no competing interest.

### Summary of Updates

The revised version corrects errors in the author information. No changes were made to the scientific content of the manuscript.

